# Bacterial membrane vesicles of *Pseudomonas Aeruginosa* activate AMPK signaling through inhibition of mitochondrial complex III

**DOI:** 10.1101/2024.06.17.599317

**Authors:** Julia Müller, Marcel Kretschmer, Elise Opitsch, Svea Holland, José Manuel Borrero-de Acuña, Dieter Jahn, Meina Neumann-Schaal, Andre Wegner

## Abstract

Bacterial membrane vesicles (BMVs) are secreted by many pathogenic bacteria and known to stimulate various host responses upon infection, thereby contributing to the pathogenicity of bacterial pathogens like *Pseudomonas aeruginosa*. While the effects of BMVs on host immune responses are well studied, little is known about their impact on cell metabolism and mitochondrial respiration. Here, we show that *P. aeruginosa* BMVs (1) reprogram cell metabolism of human lung cells, (2) negatively affect mitochondrial respiration by (3) specifically inhibiting complex III of the electron transport chain leading to (4) the activation of AMP-activated protein kinase (AMPK) signaling which in turn results in (5) AMPK-dependent inhibition of global protein synthesis.

## INTRODUCTION

The opportunistic pathogen *Pseudomonas aeruginosa* causes milder local to severe systemic infections. These infections, caused by often multi-antibiotic resistant bacteria, are particularly prevalent in immunocompromised patients, emphasizing its importance as a significant health concern in global healthcare [1, 2]. During infection, *P. aeruginosa* secretes bacterial membrane vesicles (BMVs) containing metabolites, nucleic acids and proteins, including virulence factors, which are delivered to the host cell during infection and play a pivotal role in its pathogenicity [3–5]. BMVs are well known for triggering host immune responses, capable of inducing the release of pro-inflammatory cytokines like interleukin-8 [6, 7]. Notably, BMVs can also activate the AMP-activated protein kinase (AMPK) within the host cell, leading to autophagy induction [8]. However, it remains unclear how the activation of AMPK in response to BMVs is regulated.

Given that AMPK serves as a metabolic sensor, activated by mitochondrial dysfunction [9, 10], and considering evidence of BMVs inhibiting mitochondrial activity in macrophages [11], it is conceivable that mitochondrial dysfunction might be the primary event facilitating BMV-induced AMPK activation. Some bacterial pathogens are known to affect mitochondrial function by specifically inhibiting protein complexes of the electron transport chain (ETC) which is essential for oxidative phosphorylation and ATP generation [12]. For *P. aeruginosa*, extracellular secreted factors like 1-hydroxyphenazine, pyocyanin and exotoxin A have been described as possible inhibitors of the ETC leading to reduced mitochondrial respiration [13–16]. However, the functional basis of mitochondrial dysfunction caused by BMVs is still unknown.

Here, we show that treatment of human lung cells with BMVs isolated from the pathogenic *P. aeruginosa* strain PA14 lead to metabolic reprogramming and impaired mitochondrial respiration by specifically inhibiting complex III of the ETC, resulting in mitochondrial dysfunction. Moreover, this event activates AMPK signaling, leading to AMPK-dependent inhibition of global protein synthesis in the host cell.

## METHODS

### Bacterial strains and isolation of BMVs

The bacterial strains *Pseudomonas aeruginosa* PA14 [17] and *Pseudomonas putida* KT2440 [18] were used in this study. Both strains were cultivated in lysogeny broth (LB) (Roth, X968.2) in flasks with baffles. For vesicle isolation, bacterial cultures were inoculated with an optical density measured at a wavelength of 600 nm (OD_600_) of 0.05 and grown at 37 °C (*P. aeruginosa*) or 30 °C (*P. putida*) at 160 rpm until they reached the early-stationary phase of growth. BMVs were isolated from culture media after removing bacterial cells by centrifugation at 4 °C and 8,000 x g for 30 min and filtration through 0.22 µM PES filters (Corning, 431097). Supernatants were concentrated by ultrafiltration, using Vivaspin 20 PES ultrafiltration units with a molecular weight cut-off (MWCO) of 100 kDa (Sartorius, VS2041). BMVs were isolated by ultracentrifugation at 4 °C and 150,000 x g for 2 h. Isolated vesicles were resuspended in 1X PBS (Gibco™, 18912-014) and sterile filtered. BMVs were quantified by using the membrane lipid dye FM4-64 (Invitrogen, T13320) to calculate the vesicle load per µL (VL/µL) as described before [19]. Vesicle samples were stored at 4 °C.

### Cell culture

The cell lines A549 (DSMZ, ACC 107), HCC44 (DSMZ, ACC 534) and HBEpC (PromoCell, C-12640) were used in this study. A549 cells were cultivated in DMEM medium (Gibco™, 11965-092) containing 25 mM glucose and 4 mM glutamine. HCC44 cells were cultivated in RPMI 1640 medium (Gibco™, 21875-034) containing 11 mM glucose and 2 mM glutamine. Both growth media were supplemented with 10 % FBS (Bio&SELL, FBS.SAM.0500). HBEpC cells were cultivated in Airway Epithelial Cell Growth Medium supplemented with Growth Medium SupplementMix (PromoCell, C-21060). All cell lines were incubated in a humidified atmosphere with 5 % CO_2_ at 37 °C. Cell detachment was performed with 0.05 % trypsin-EDTA (Gibco™, 25300-054) for A549 and HCC44 cells or accutase solution (Sigma-Aldrich, A6964) for HBEpC cells.

### Analysis of cell growth and viability

To analyze cell growth of the proliferating cell lines A549 and HCC44, cell confluence was measured in 6-well plates at 37 °C with a microplate reader (Tecan Spark^®^). For the analysis of cell viability of non-proliferating HBEpC cells, the PrestoBlue™Cell Viability Reagent (Invitrogen™, A13261) was used according to the user manual. Fluorescence signals were measured by using an excitation wavelength of 560 nm and emission wavelength of 590 nm.

### Stable isotope labeling

For stable isotope labeling, 150,000 cells/well (A549), 100,000 cells/well (HCC44) or 70,000 cells/well (HBEpC) were seeded in 6-well plates (Greiner Bio-One, 657160) and incubated at 37 °C and 5 % CO_2_ overnight. The next day, cells were washed with 1X PBS and treated as indicated in the figure legends in the following labeling media: A549 cells were cultivated in DMEM medium (Gibco™, A14430-01) supplemented with 25 mM unlabeled glucose or [U-^13^C_6_]-glucose and 4 mM [U-^13^C_5_]- glutamine or unlabeled glutamine and 10 % dFBS. HCC44 cells were cultivated in SILAC RPMI 1640 medium (Gibco™, A24942-01) supplemented with 11.1 mM unlabeled glucose or [U-^13^C_6_]-glucose, 2.05 mM [U-^13^C_5_]-glutamine or unlabeled glutamine, 0.22 mM lysine, 1.15 mM arginine and 10 % dFBS. HBEpC cells were cultivated in DMEM medium (Gibco™, A14430-01) supplemented with 25 mM unlabeled glucose or [U-^13^C_6_]-glucose, 4 mM [U-^13^C_5_]-glutamine or unlabeled glutamine and 2 % Growth Medium SupplementMix (PromoCell, C-39165).

### Metabolite extraction and GC/MS analysis

For GC/MS analysis of intracellular metabolites, 150,000 cells/well (A549) or 70,000 cells/well (HBEpC) were seeded in 6-well plates (Greiner Bio-One, 657160) and incubated at 37 °C and 5 % CO_2_ overnight. The next day, cells were washed with 1X PBS and treated as indicated in the figure legends. Treated cells were extracted as described by Sapcariu et al. [20]. Briefly, cells were washed with 2 mL/well 0.9 % sodium chloride and 400 µL/well methanol (-20 °C) was added. Then, 400 µL/well water (4 °C), containing 1 µg/mL glutaric acid-d_6_ as an internal standard, was added. Cells were mechanically detached, the cell suspension transferred into 400 µL chloroform (-20 °C) and mixed at 4 °C and 1,400 rpm for 20 min. To separate polar and non-polar phases, samples were centrifuged at 4 °C and 17,000 x g for 5 min. The upper polar phase was transferred in GC/MS vials and dried in a speedvac at 4 °C overnight. Dried samples were derivatized with 15 µL of 2 % (w/v) methoxyamine-hydrochloride solved in pyridine by shaking at 40 °C for 90 min and additional15 µL of *N*-methyl-*N*-(trimethylsilyl)trifluoroacetamide (MSTFA) by shaking at 40 °C for 30 min or *N*-*tert* -butyldimethylsilyl-*N*-methyltrifluoroacetamide (MTBSTFA) by shaking at 55 °C for 60 min. The derivatized samples (1 µL) were injected into a SSL injector in splitless mode and heated up to 270 °C. GC/MS measurements were performed with an Agilent Technologies 7890B GC system including a 30 m Phenomenex ZB-35 and 5 m Duraguard capillary column, connected to an Agilent Technologies 5977B MSD, under electron ionization at 70 eV. The MS source temperature was held at 230 °C and the quadrupole temperature at 150 °C. Helium was used as a carrier gas with a flow rate of 1 mL/min. The temperature profile of the GC oven depended on the used measuring method. For the measurement of MTBSTFA-derivatized samples, the GC oven temperature was held at 100 °C for 2 min, then increased up to 300 °C at 10 °C/min and held at 300 °C for 4 min. For the measurement of MSTFA-derivatized samples, the GC oven temperature was held at 80 °C for 6 min, then increased up to 300 °C at 6 °C/min, held at 300 °C for 10 min, raised to 325 °C at 10 °C/min and held at 325 °C for 4 min. The total abundances of metabolites and distributions of mass isotopomers were calculated by the integration of mass fragments and corrected for natural isotope abundances by using the software MetaboliteDetector as previously described [21].

### Quantitative PCR analysis

RNA was isolated from the interphase of extracted cells as described before [20] by using the NucleoSpin^®^ RNA kit (MACHEREY-NAGEL, 740955.50). Isolated RNA was converted to cDNA by using the High-Capacity cDNA Reverse Transcription Kit (Applied Biosystems™, 4368813). RNA and cDNA concentrations were measured with a microplate reader (Tecan Spark^®^). For quantitative PCR (qPCR) analysis, TaqMan™gene expression assays (Applied Biosystems™) for the housekeeping gene 18S (Hs99999901_s1) and target genes HMGCR (Hs00168352_m1) and SREBF2 (Hs01081784_m1) were used, together with the iTaq™Universal Probes Supermix (Bio-Rad, 1725132) and the QuantStudio™5 Real-Time PCR system (Applied Biosystems™). Data was analyzed with the QuantStudio™design and analysis software.

### Analysis of uptake and secretion rates

Glucose uptake and lactate secretion were measured by using the YSI 2950 biochemistry analyzer. To that end, medium samples were collected before metabolite extraction, centrifuged at 4 °C and 17,000 xg for 5 min and 200 µL of each sample was transferred in a 96-well microplate. For calibration, a standard solution containing 28 mM glucose and 15 mM lactate was measured in a dilution series of 5, 10, 20, 40, 60, 80 and 100 %.

### Protein extraction and immunoblotting

For protein analysis, A549 (1,000,000 cells), HCC44 (500,000 cells) and HBEpC cells (500,000 cells) were plated in 10 cm (A549 and HCC44 cells) or 6 cm (HBEpC cells) cell culture dishes (Greiner Bio-One) in growth medium and incubated at 37 °C and 5 % CO_2_ overnight. The next day, cells were treated as indicated in the figure legends. After treatment, cells were lysed with the M-PER extraction reagent (Thermo Scientific, 78501) containing 1X Halt protease and phosphatase inhibitors (Thermo Scientific, 78441), mixed at 4 °C and 1,400 rpm for 10 min and centrifuged at 4 °C and 14,000 xg for 10 min. Supernatants containing protein were stored at -20 °C. Protein quantification was performed by using the Pierce BCA protein assay kit (Thermo Scientific, 23227) according to the user manual. Samples were loaded with a 5X Laemmli buffer, incubated at 95 °C for 5 min and centrifuged at 16,000 xg for 1 min. 25 µg of total protein was separated on 4-20 % precast SDS-PAGE gels (Bio-Rad). Depending on the protein size, a prestained protein marker covering the range of 10-180 kDa (Thermo Scientific, 26616) or 43-315 kDa (Cell Signaling Technology, 12949) was used. Proteins were transferred onto a 0.45 µm PVDF membrane (Carl Roth, T830.1) by using the Trans-Blot^®^ SD Semi-Dry Transfer Cell (Bio-Rad). The membrane was blocked with 5 % BSA (w/v) (Biomol, 01400.100) in Tris-buffered saline with 0.1 % Tween 20 (TBS-T) for 1 h. Primary antibodies diluted in 1X TBS-T with 5 % BSA (Table 1) were added to the membranes and incubated at room temperature for 1 h. Membranes were then incubated with secondary antibodies diluted in 1X TBS-T with 5 % BSA (Table 2) at room temperature for 1 h. For signal detection, the Immobilon Classico Western HRP substrate (Millipore, WBLUC0500) was used and imaged with a Bio-Rad ChemiDoc imaging system. Band intensities were analyzed by using the ImageJ 1.53k software.

**Table 1:** Primary antibodies.

| Primary antibody | Catalog no. | Company | Dilution |
| --- | --- | --- | --- |
| Mouse monoclonal anti-β-actin | A5441 | Merck | 1:5,000 |
| Rabbit polyclonal anti-ACC1 | 21923-1-AP | Proteintech | 1:4,000 |
| Rabbit polyclonal anti-phospho-ACC1 (Ser79) | 29119-1-AP | Proteintech | 1:1,000 |
| Rabbit polyclonal anti-eEF2K | 13510-1-AP | Proteintech | 1:1,000 |
| Rabbit polyclonal anti-phospho-eEF2K (Ser366) | 29032-1-AP | Proteintech | 1:2,000 |
| Rabbit polyclonal anti-eEF2 | 20107-1-AP | Proteintech | 1:10,000 |
| Rabbit polyclonal anti-phospho-eEF2 (Thr56) | E-AB-51049 | Elabscience | 1:1,000 |

**Table 2:** Secondary antibodies.

| Primary antibody | Catalog no. | Company | Dilution |
| --- | --- | --- | --- |
| Goat anti-mouse, HRP-conjugated | R-05071-500 | Advansta | 1:20,000 |
| Goat anti-rabbit, HRP-conjugated | R-05072-500 | Advansta | 1:20,000 |

### Analysis of global protein synthesis

For the analysis of global protein synthesis, 30,000 cells/well (A549) were seeded in black 96-well microplates (Greiner Bio-One, 655090) in growth medium and incubated at 37 °C and 5 % CO_2_ overnight. The next day, treatments were performed as indicated in the figure legends. Protein synthesis of treated cells was measured by using the Protein Synthesis Assay Kit (Cayman Chemical, 601100) according to the user manual.

### Measurement of interleukin-8 secretion

To analyze IL-8 secretion, 150,000 cells/well (A549) or 100,000 cells/well (HBEpC) were seeded in 6-well plates in growth medium and incubated at 37 °C and 5 % CO_2_ overnight. The next day, cells were treated with 25 VL/mL isolated PA14 WT vesicles and incubated at 37 °C and 5 % CO_2_ for 24 h. After treatment, medium was collected and centrifuged at 4 °C and 17,000 x g for 5 min to remove cell debris. IL-8 concentration in medium samples was measured by using the ELISA MAX Deluxe Set Human IL-8 (BioLegend, 431504) according to the user manual.

### RNA-Sequencing analysis

Cells were seeded in 6-well plates and incubated at 37 °C and 5 % CO_2_ overnight. The next day, cells were washed with 1X PBS, treated with BMVs and incubated at 37 °C and 5 % CO_2_ for 24 h. RNA isolation was performed by using the NucleoSpin^®^ RNA kit (MACHEREY-NAGEL, 740955.50) as described above. cDNA synthesis was performed and libraries were constructed using NEBNext Ultra II Directional RNA Library Prep Kit for Illumina at the Genome Analytics at Helmholtz Centre for Infection Research. Libraries were sequenced using a NovaSeq 6000 instrument (Illumina) generating 50-bp reads in paired-end mode. Reads mapping and differential expression analysis were performed using the galaxy platform (https://usegalaxy.org/) [22]. Reads were mapped to the human genome hg19 with Hisat2 [23]. EdgeR was used to identify differential expression and calculate the P values with an exact test based on the dispersion generated by the quantile-adjusted conditional maximum likelihood (qCML) method [24].

### Respirometry

Cell respiration was measured by using an Agilent Seahorse XFe96 Analyzer together with Seahorse XFe96 Extracellular Flux Assay Kits. Cells were seeded in Seahorse XF96 V3 PS Cell Culture Microplates (Agilent, 101085-004) and cultured in growth medium at 37 °C and 5 % CO_2_ overnight. At the same time, the sensor cartridge (Agilent, 103792-100) was incubated with sterile water together with the Seahorse XF Calibrant Solution (Agilent, 100840-000) at 37 °C and 0 % CO_2_. The next day, the medium was replaced with Seahorse XF DMEM medium, pH 7.4 (Agilent, 103575-100) supplemented with 10 mM glucose, 2 mM glutamine and 10 % dFBs and incubated at 37 °C and 0 % CO_2_ for 60 min. For sensor cartridge calibration, the water was replaced with Seahorse XF Calibrant Solution and incubated at 37 °C and 0 % CO_2_ for 45 min. To assess mitochondrial function, the oxygen consumption rate (pmol O_2_/min) was measured and normalized to basal respiration. Mitochondrial respiratory chain deficiencies were analyzed based on the study of Jaber et al. [25].

## RESULTS AND DISCUSSION

### *P. aeruginosa* BMVs inhibit proliferation and viability of human lung cells

BMVs from several bacterial species are reported to inhibit the proliferation of different host cell types [19, 26–28]. Thus, we aimed to explore the impact of *P. aeruginosa* BMVs on the proliferation of human lung cells in initial experiments. To this end, we treated A549 and HCC44 lung cancer cells with BMVs isolated from the pathogenic *P. aeruginosa* strain PA14 and monitored their cell confluence over 72 hours. Moreover, we analyzed the effects of PA14 BMVs on the viability of primary bronchial epithelial cells (HBEpC) by fluorescently labeling living cells, as these cells are non-proliferating. For all cell lines, we observed a significant decrease in confluence (Figure 1A) or cell viability (Figure S1A) after vesicle treatment, highlighting the pathogenic potential of PA14 BMVs. In contrast, treatment with BMVs isolated from the non-pathogenic strain *Pseudomonas putida* KT2440 resulted in a significantly weaker effect (Figure S1A and B). This suggests that the specific cargo of the pathogenic PA14 BMVs was responsible for the host reaction, rather than BMVs in general. However, the precise molecular mechanism of the observed anti-proliferative effect remains unknown.

**Figure 1:**
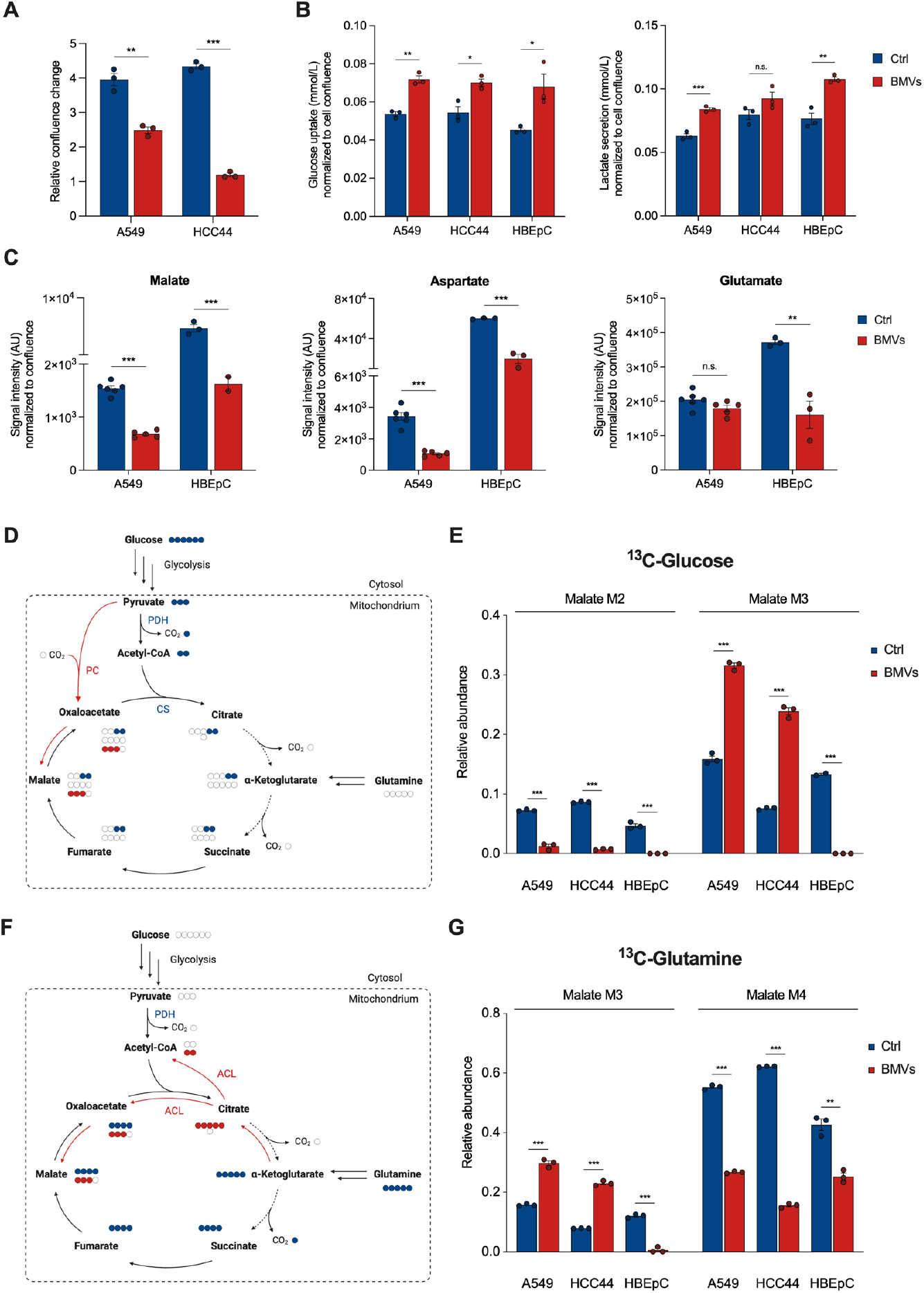
Metabolic reprogramming in host cells after PA14 BMV treatment. **(A)** Confluence of A549 and HCC44 cells after treatment with 25 VL/mL BMVs for 72 h. Data were obtained from 3 biological replicates. **(B)** Glucose uptake and lactate secretion of A549, HCC44 and HBEpC cells after treatment with 25 VL/mL BMVs for 24 h. Data were obtained from 3 biological replicates. **(C)** Signal intensities (AU) of malate, aspartate and glutamate in A549 and HBEpC cells after vesicle treatment (25 VL/mL for 24 h). Data were obtained from 5 - 6 (A549) or 2 - 3 (HBEpC) biological replicates and normalized to cell confluence. **(D)** Schematic overview of the incorporation of a [U-^13^C6]-glucose tracer into metabolites of the TCA cycle. Created with BioRender.com. **(E)** Malate MIDs after [U-^13^C6]-glucose labeling of BMV-treated cells. A549 and HCC44 cells were treated for 24 h and HBEpC cells for 6 h with 25 VL/mL. Data were obtained from 2 or 3 biological replicates. **(F)** Schematic overview of the incorporation of a [U-^13^C5]-glutamine tracer into metabolites of the TCA cycle. Created with BioRender.com. **(G)** Malate MIDs after [U-^13^C5]-glutamine labeling of BMV-treated cells. A549 and HCC44 cells were treated for 24 h and HBEpC cells for 6 h with 25 VL/mL. Data were obtained from 2 or 3 biological replicates. All bar plots in this Figure are depicted as mean SEM. All significance niveaus were determined by Student’s t-test (n.s. = not significant, * = p < 0.05, ** = p < 0.01, *** = p < 0.001).

### *P. aeruginosa* BMVs induce rapid metabolic reprogramming by modulating TCA cycle activity

BMVs have been recognized for modulating the host immune response during *P. aeruginosa* infection [6, 7]. However, the underlying molecular dynamics remain elusive. Given the close link between cellular functions and cell metabolism, investigating the metabolic responses of the host cell induced by PA14 BMVs would be revealing for the understanding of the host-pathogen interaction of *P. aeruginosa*. Recently, we developed a method for the isolation and quantification of BMVs in order to analyze vesicle-induced metabolic changes in mammalian cell cultures [19]. To ascertain the potential metabolic effects induced by BMVs, we initially examined changes in glucose uptake and lactate secretion of vesicle-treated lung cancer cells as well as primary bronchial epithelial cells. We observed a significant increase in glucose uptake and lactate secretion for all tested cell lines (Figure 1B), indicating increased glycolytic activity. This phenomenon is known for various bacterial and viral infections, especially in immune cells, presumably to provide biosynthetic intermediates for the synthesis of nucleotides, amino acids and lipids to support host cell proliferation [29–31].

To further investigate BMV-induced metabolic shifts in host cells, we employed gas chromatography coupled with mass spectrometry (GC/MS) to analyze BMV-treated A549, HCC44 and HBEpC cells. Our results indicate broad metabolic reprogramming of both lung cancer and primary lung cells. Specifically, we observed decreased levels of TCA cycle-associated metabolites like malate, aspartate and glutamate in cells treated with PA14 BMVs (Figure 1C). Analogous to cell growth, these effects were significantly reduced when cells were treated with BMVs of the non-pathogenic *P. putida* KT2440 strain (Figure S1C). Conversely, the levels of most amino acids were increased in A549 cells after BMV treatment, presumably due to autophagy induction, as previously reported [19].

To understand the metabolic pathways contributing to these observations, we used stable isotope-assisted metabolomics. By feeding cells with [U-^13^C_6_]-glucose and [U-^13^C_5_]-glutamine during vesicle exposure, we noted significant shifts in the mass isotopomer distribution (MID) of TCA cycle-associated metabolites, exemplified by malate. As glucose can enter the TCA cycle via acetyl-CoA, the use of the [U-^13^C_6_]-glucose tracer results in the formation of M2 citrate (Figure 1D) and, for the oxidative TCA cycle flux, in M2 malate. Interestingly, the formation of M2 malate was significantly decreased in all tested cell lines after vesicle treatment (Figure 1E, p < 0.001). For A549 and HCC44 cells, we also observed an increased formation of M3 malate from [U-^13^C_6_]-glucose, involving the conversion of fully labeled pyruvate to oxaloacetate by carboxylation via pyruvate carboxylase (PC). In this step, free unlabeled CO_2_ gets incorporated into oxaloacetate resulting in M3 malate (Figure 1D). The significantly increased formation of M3 malate in A549 and HCC44 cells after BMV treatment (Figure 1E, p < 0.001) suggests a shift towards increased PC activity, presumably to compensate for the reduced oxidative TCA cycle flux. However, we did not observe this effect in HBEpC cells, indicating an inactive PC and no possibility to evade the affected flux.

Using [U-^13^C_5_]-glutamine as a substrate, active oxidative TCA cycle flux results in M4 malate (Figure 1F). After BMV treatment, we observed decreased formation of M4 malate from [U-^13^C_5_]-glutamine for all tested cell lines (Figure 1G), confirming the ^13^C-glucose results. Moreover, the formation of M3 malate significantly increased in A549 and HCC44 cells (Figure 1G, p < 0.001), explained by a shift to the reductive TCA cycle flux [32], resulting in M5 citrate that can be converted to M2 acetyl-CoA and M3 oxaloacetate via ATP-citrate lyase (ACL, Figure 1F). Similar to the glucose labeling results, we did not observe the increased formation of M3 malate in HBEpC cells (Figure 1G). Deregulation of enzymes like PC and ATP-citrate lyase, associated with the reductive TCA cycle flux, is known for various pathological conditions like cancer, but also in infection [33, 34]. Our results suggest that the shift towards a higher activity of these enzymes after BMV treatment is specific for lung cancer cell metabolism and does not occur in primary lung cells. To investigate whether these effects are caused by specific virulence agents of PA14 BMVs or vesicles in general, we also analyzed the effects of BMVs isolated from the non-virulent *P. putida* KT2440 strain. We observed much smaller effects when cells were treated with these vesicles which further suggests that the effects are due to specific factors contained in BMVs isolated from the pathogenic *P. aeruginosa* PA14 strain (Figure S1C, D, and E).

To analyze how fast the observed metabolic changes appear after BMV treatment, we analyzed the effects after different treatment times (15, 60, and 240 minutes). We observed rapid metabolic reprogramming following BMV treatment, with decreased M2 and increased M3 malate already observable after 15 minutes in A549 cells (Figure S2A). Moreover, the cellular malate levels were reduced by 20 % at this time (Figure S2B).

Our results indicate a rapid PA14 vesicle-driven impact on cell metabolism, evidenced by the decreased oxidative TCA cycle flux following BMV treatment. The swift onset of this effect suggests that the observed metabolic changes are not due to prior cellular alterations, such as changes at gene expression level. Instead, this indicates the presence of a vesicle-associated factor that triggers an immediate effect on cellular respiration.

### *P. aeruginosa* BMVs impair mitochondrial respiration of human lung cells by inhibiting electron transport chain complex III

The aforementioned metabolic changes indicate a reduced oxidative TCA cycle flux after BMV treatment, resulting in decreased generation of NADH and FADH_2_, which are essential for mitochondrial respiration and ATP production. Deo et al. have previously demonstrated the inhibitory effects of *P. aeruginosa* BMVs on mitochondrial activity in macrophages, leading to mitochondrial apoptosis and inflammation [11]. Consistent with these results, we observed a significantly higher NADH/NAD^+^ ratio, and decreased ATP levels in A549 and HCC44 cells after BMV treatment (Figure 2A and B).

**Figure 2:**
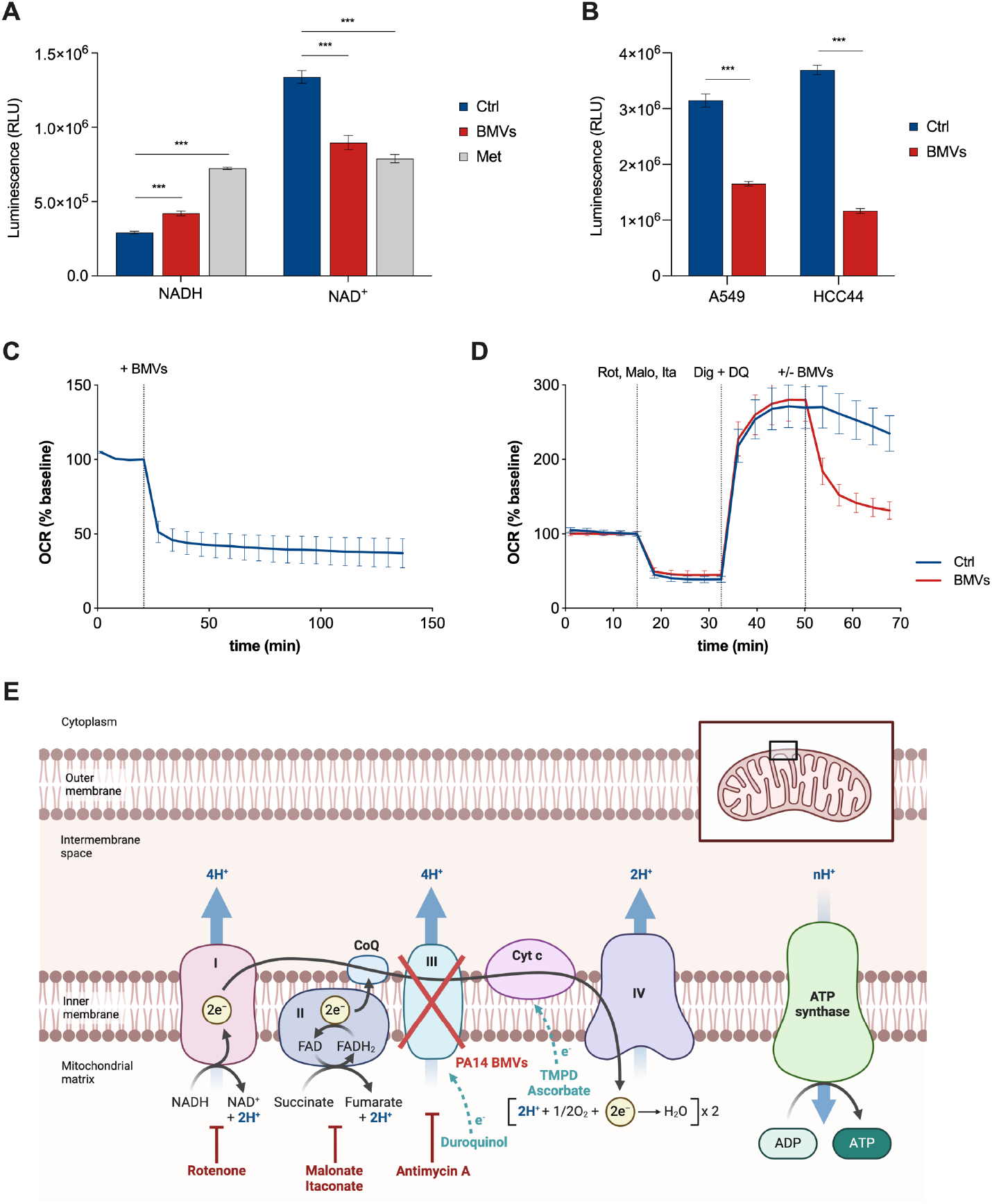
PA14 BMVs induce mitochondrial dysfunction in host cells. **(A)** NADH and NAD^+^ levels of A549 cells after treatment with 25 VL/mL BMVs and 1 mM metformin (Met) for 24 h measured as luminescence intensities (RLU). Data were obtained from 15 biological replicates. **(B)** ATP levels of A549 and HCC44 cells after BMV treatment (25 VL/mL for 24 h) measured as luminescence intensities (RLU). Data were obtained from 10 biological replicates. **(C)** OCR of A549 cells after BMV treatment. 25 VL/mL BMVs were added at the depicted time point. Data were obtained from 15 biological replicates. **(D)** Respirometry test for complex III activity of the ETC of A459 cells after BMV treatment (25 VL/mL). Isolation of complex III was performed by the addition of 2 µM rotenone (Rot), 40 µM malonate (Malo), 2 mM itaconate (Ita), 25 µg/mL digitonin (Dig) and 1 mM duroquinol (DQ) at the depicted time points. Data were obtained from 30 biological replicates. All OCR values in this Figure were normalized to the baseline and depicted as mean SEM. **(E)** Location of electron transport deficiencies by functionally isolating Cytochrome *c* and the complexes of the ETC to test their activity after vesicle treatment. Rotenone, malonate and itaconate or Antimycin A are added to inhibit complex I, II or III which interrupts the electron transfer to the following complexes. The electron transport can be restored by the addition of the electron donor duroquinol for complex III or TMPD and its reducing agent ascorbate for Cytochrome *c*. PA14 BMVs are able to specifically inhibit complex III of the ETC. All bar plots in this Figure are depicted as mean *±* SEM. Significance niveaus were determined by Student’s t-test (*** = p < 0.001).

To validate, if *P. aeruginosa* BMVs alter mitochondrial respiration in human lung cells, we analyzed changes of cellular respiration by measuring the oxygen consumption rate (OCR) in BMV-infected A549 cells. We observed an decreased OCR immediately after vesicle treatment (Figure 2C), indicating a rapid effect on mitochondrial respiration (< 7 minutes) which might explain the swift metabolic adaptation discussed above.

Mitochondrial respiration relies on oxidative phosphorylation, which requires the electron transport chain (ETC) located in the inner mitochondrial membrane and composed of complexes I, II, III and IV, as well as proteins like Cytochrome *c*. It is well known that some bacterial pathogens are able to inhibit complexes of the ETC [12]. For *P. aeruginosa*, the extracellular secreted factors exotoxin A, 1-hydroxyphenazine and pyocyanin have been reported to affect mitochondrial respiration, presumably by disrupting the ETC [13–16].

To determine whether *P. aeruginosa* BMVs specifically impair one of the ETC complexes, leading to the observed respiratory deficit, we followed the protocol by Jaber et al. [25]. This protocol involves a stepwise series of experiments to map the location of electron transport deficiencies, beginning with Cytochrome *c*. If BMVs cause a Cytochrome *c* deficit, then adding exogenous Cytochrome *c* should rescue respiration. Because Cytochrome *c* cannot cross the plasma membrane, we selectively permeabilized the plasma membrane using digitonin. However, we did not observe a rescue effect in the OCR of BMV-treated A549 cells supplemented with Cytochrome *c* (Figure S3A), suggesting that a deficit in another component of the ETC is limiting respiration.

After excluding Cytochrome *c* deficiency, we functionally isolated the individual complexes of the electron transport chain by selectively limiting electron entry to each complex and determining whether the deficiency can still be observed. Since complexes I to IV function in series, a dysfunctional complex would affect subsequent complexes. For this reason, we started with complex IV.

To functionally isolate complex IV, we inhibited complex III with Antimycin A (Figure 2E), which interrupts electron transfer to complex IV, leading to a decreased OCR (Figure S3B). We then added N,N,N’,N’-tetramethyl-p-phenylenediamine (TMPD) and its reducing agent ascorbate to restore electron transfer to complex IV, bypassing complex III (Figure 2E). We observed that complex IV activity was not affected by BMVs (Figure S3B), leaving complexes I to III as potential targets.

Next, we analyzed complex III activity by inhibiting complex I with rotenone and complex II with malonate and itaconate (Figure 2E) [35, 36]. To ensure electron transfer to complex III, we added the electron donor duroquinol, which recovers respiration by feeding electrons into complex III (Figure 2E). Interestingly, we observed impaired complex III activity immediately after vesicle addition, indicating specific inhibition of this complex by PA14 BMVs (Figure 2D). Moreover, we observed impairment of complexes I and II, presumably as a consequence of complex III inhibition (Figure S3C and D).

Overall, these findings suggest that PA14 BMVs specifically target complex III of the ETC, leading to a cascade of inhibition that affects complexes I and II, thereby impairing overall mitochondrial respiration.

### *P. aeruginosa* BMVs suppress cholesterol biosynthesis and global protein synthesis while inducing inflammation

Since BMVs are able to induce drastic effects on host cell metabolism, we assumed that other general cellular processes would also be impaired. To gain an overview of altered cellular functions, we performed RNA-Sequencing analysis followed by pathway enrichment analysis. We identified the suppression of cholesterol biosynthesis as the most affected pathway in A549 cells treated with PA14 BMVs (Figure S4). Suppression of cholesterol biosynthesis is well-documented in viral infections [37], but less explored in bacterial infections. Strikingly, we observed a downregulation of all genes associated with the cholesterol biosynthesis pathway (Figure 3A). For further validation, we analyzed the expression of selected downregulated genes by using quantitative PCR (Figure 3B). We observed a significantly decreased expression of 3-hydroxy-3-methylglutaryl-coenzyme A reductase (HMGCR, p < 0.001) and sterol regulatory element binding transcription factor 2 (SREBF2, p < 0.05) after 24 h of BMV exposure. HMGCR is the rate-limiting enzyme of the cholesterol synthesis pathway [38], and its expression can be activated by SREBF2 which regulates various key enzymes associated with cholesterol and fatty acid synthesis [39, 40].

**Figure 3:**
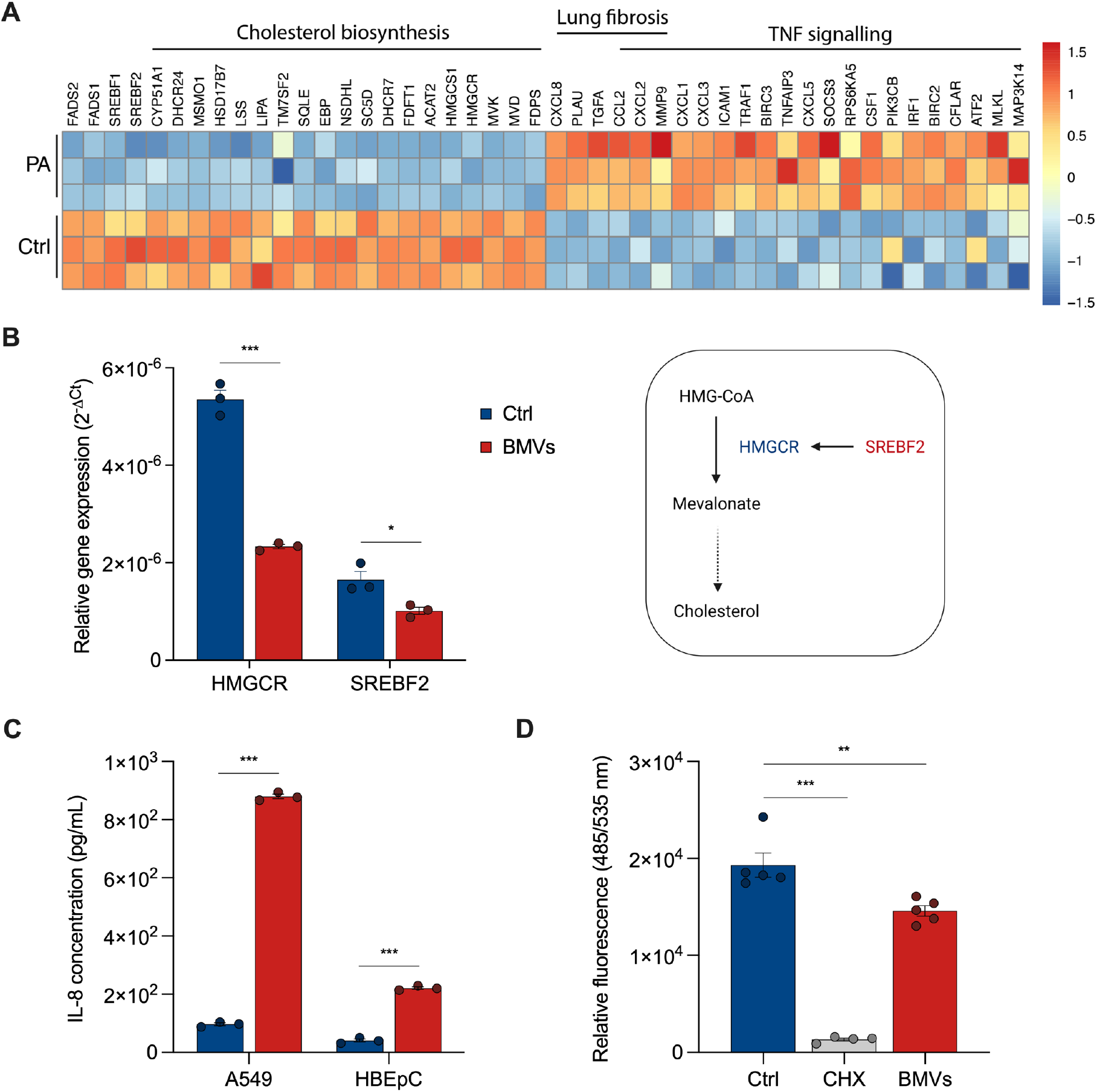
PA14 BMVs affect transcription and translation of the host cell. **(A)** RNA-Sequencing analysis of A549 cells treated with PA14 BMVs for 24 h. **(B)** Relative gene expression of HMGCR and SREBF2 in A549 cells after treatment with 25 VL/mL BMVs for 24 h. Data were obtained from 3 biological replicates. **(C)** IL-8 secretion of A549 and HBEpC cells after BMV treatment (25 VL/mL for 24 h). Data were obtained from 3 biological replicates. **(D)** Global protein synthesis of A549 cells after treatment with 50 µg/mL cycloheximide (CHX) and 25 VL/mL BMVs for 30 min (+ 30 min pre-treatment). Data were obtained from 4 or 5 biological replicates. All bar plots in this Figure are depicted as mean SEM. All significance niveaus were determined by Student’s t-test (* = p < 0.05, ** = p < 0.01, *** = p < 0.001).

We also observed an upregulation of markers associated with inflammation (TNF signaling) and disease (lung fibrosis) after vesicle treatment (Figure 3A). For example, we determined a higher expression of interleukin-8 (IL-8, *CXCL8*) which plays a key role in acute inflammation [41]. Bauman and Kuehn have previously shown that vesicles isolated from the *P. aeruginosa* PAO1 strain induce IL-8 activation in lung epithelial cells [6]. To confirm this result using PA14 BMVs, we exposed lung cancer cells (A549) and primary lung cells (HBEpC) to BMVs for 24 h and measured their IL-8 secretion (Figure 3C). We observed a significantly increased IL-8 secretion for both cell lines (p < 0.001). The stronger effect in A549 cells could be explained by the general importance of IL-8 in cancer progression [42].

Next, we analyzed whether vesicles affect not only transcription, but also translation in the host cell, since there are known bacterial regulators of protein translation, such as the exotoxin A of *P. aeruginosa* [43]. Moreover, inhibition of protein synthesis by BMVs has already been described in macrophages due to mitochondrial stress [11]. To confirm similar changes in human lung cells, we labeled newly translated proteins of A549 cells with a fluorescent dye and analyzed their signal intensities after treatment with the translation inhibitor cycloheximide (CHX) and PA14 BMVs. We observed a significantly decreased fluorescence signal for both treatments already after 30 minutes, indicating the inhibition of global protein synthesis by PA14 BMVs in A549 cells (Figure 3D).

### *P. aeruginosa* BMVs activate AMPK signaling through mitochondrial dysfunction, leading to global protein synthesis inhibition

We showed that PA14 BMVs are able to affect general cellular functions of the host cell, suggesting the activation of a global signaling pathway inside the cell. The AMP-activated protein kinase (AMPK) is known to function as a metabolic sensor that can be activated e.g. by mitochondrial dysfunction, sensing decreased ATP levels caused by ETC inhibition [44]. Moreover, mitochondria-localized AMPK is known to enable mitochondrial function [45]. Losier et al. demonstrated that AMPK is stimulated by the detection of BMVs during infection with the pathogen *Salmonella enterica* serovar Typhimurium, resulting in autophagy induction [8]. Interestingly, AMPK activation is known to downregulate HMGCR which in turn suppresses cholesterol synthesis [46], aligning with our result that HMGCR gene expression is decreased after BMV treatment (Figure 3A and B).

To confirm whether AMPK signaling is activated by PA14 BMVs, we treated A549 cells with *P. aeruginosa* BMVs and analyzed the activation of the AMPK target enzyme acetyl-CoA carboxylase 1 (ACC1, Figure 4A) by determining its phosphorylation via western blot analysis. When phosphorylated, ACC1 inhibits the conversion of acetyl-CoA to malonyl-CoA, which is needed for fatty acid synthesis [47]. We observed a general decrease in ACC1 protein levels after vesicle treatment, while the p-ACC1/ACC1 ratio increased in a concentration-dependent manner, indicating the activation of AMPK by PA14 BMVs (Figure 4B). This result also suggests a regulation of ACC1 protein translation by BMVs, aligning with our observation that PA14 BMVs inhibit global protein synthesis in the host cell (Figure 3D). Since AMPK signaling can regulate protein translation via the eukaryotic elongation factor 2 (eEF2), we analyzed its activation by western blot analysis and observed a highly increased p-eEF2/eEF2 ratio after BMV treatment (Figure 4C). As eEF2 acts as a negative regulator of protein synthesis [48], its activation indicates an inhibition of global protein synthesis by PA14 BMVs. We observed the activation of eEF2 in lung cancer cells (Figure 4C and D) as well as in primary lung cells (Figure 4E), with a higher phosphorylation increase in cancer cells. Taken together, our results suggest that PA14 BMVs inhibit global protein synthesis in both lung cancer and primary lung cells, with a more pronounced effect in cancer cells.

**Figure 4:**
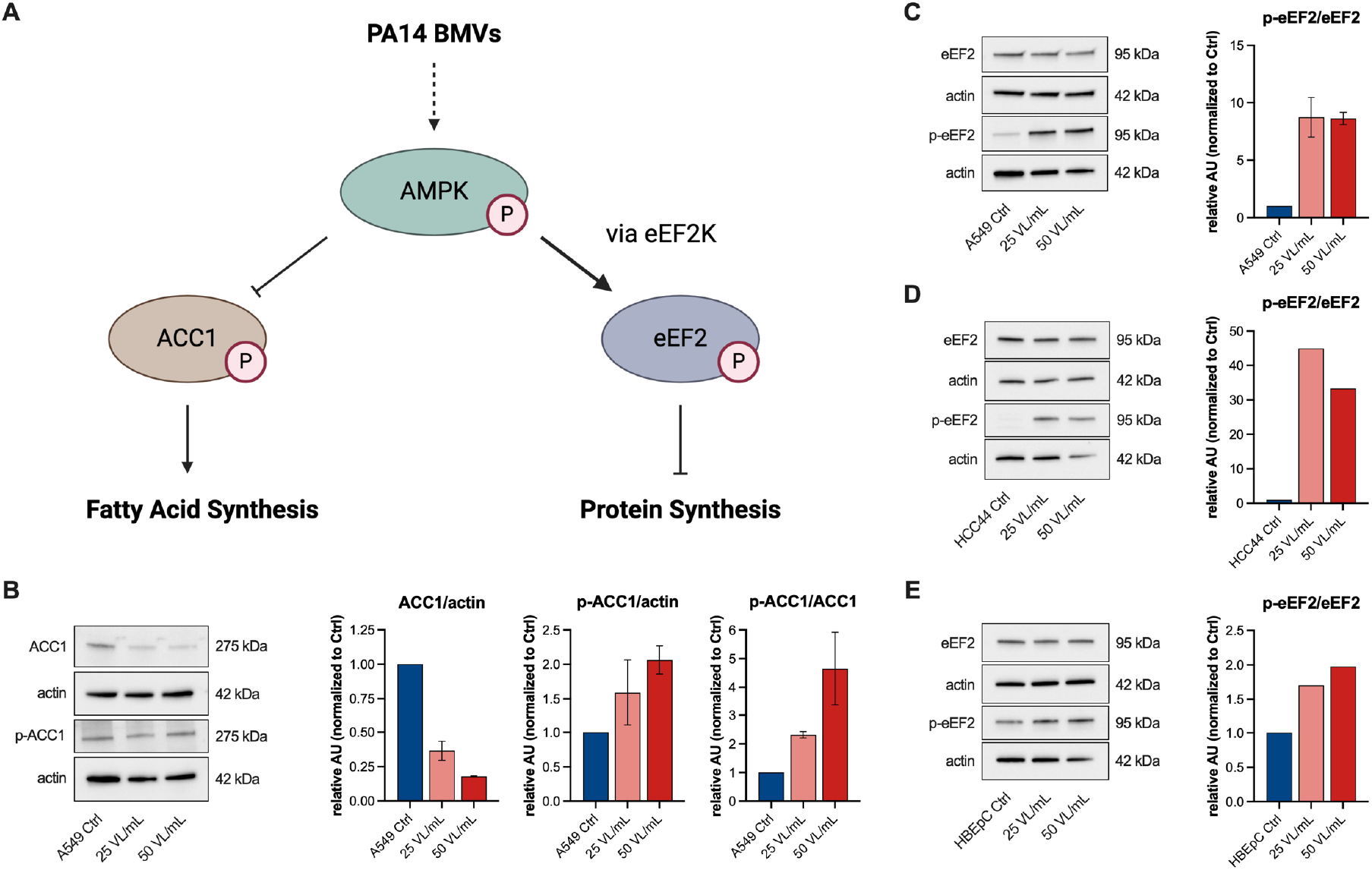
Activation of AMPK signaling in host cells by PA14 BMVs. **(A)** Overview of the AMPK signaling targets ACC1 and eEF2 and their activation. **(B)** WB analysis of p-ACC1 (Ser79) in A549 cells after vesicle treatment (25 VL/mL and 50 VL/mL) for 24 h. Data were obtained from 2 independent experiments. **(C)** WB analysis of p-eEF2 (Thr56) in A549 cells after vesicle treatment (25 VL/mL and 50 VL/mL) for 24 h. Data were obtained from 2 independent experiments. **(D)** WB analysis of p-eEF2 (Thr56) in HCC44 cells after vesicle treatment (25 VL/mL and 50 VL/mL) for 24 h. **(E)** WB analysis of p-eEF2 (Thr56) in HBEpC cells after vesicle treatment (25 VL/mL and 50 VL/mL) for 24 h. All bar plots in this Figure are depicted as mean (*±* SEM).

Since it has also been shown that BMVs of *Salmonella enterica* serovar Typhimurium inhibit mTOR signaling [8], and given that eEF2 can also be regulated by mTOR, we analyzed the activity of the upstream kinase eEF2K, using an antibody for the mTOR-specific phosphorylation site Ser366 [48, 49]. However, we did not observe any change in phosphorylation at this site (Figure S5), suggesting that the vesicle-driven effects on protein synthesis are regulated independently of mTOR. To confirm that translation is inhibited via AMPK signaling, we treated BMV-infected A549 cells with the AMPK inhibitor Compound C (CC) and analyzed the activity of ACC1 and eEF2. We discovered that AMPK inhibition suppressed the vesicle-mediated activation of both targets (Figure 5A and B), connecting the inhibition of protein synthesis by BMVs to the activation of AMPK signaling.

**Figure 5:**
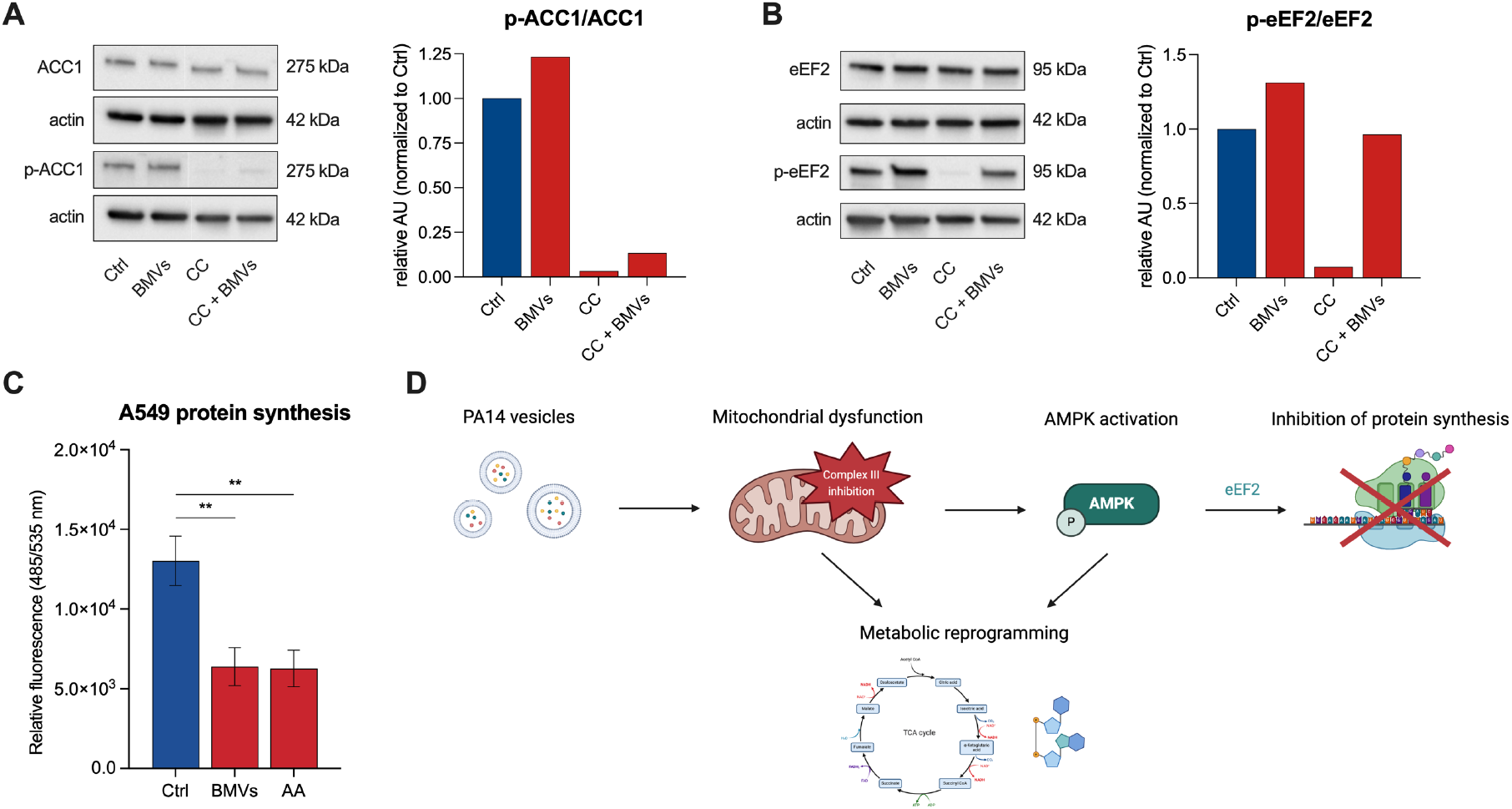
AMPK-mediated inhibition of global protein synthesis through ETC inhibition after PA14 BMV treatment. **(A)** WB analysis of p-ACC1 (Ser79) in A549 cells after treatment with 25 VL/mL BMVs and 10 µM Compound C (CC) for 24 h. **(B)** WB analysis of p-eEF2 (Thr56) in A549 cells after treatment with 25 VL/mL BMVs and 10 µM Compound C (CC) for 24 h. **(C)** Global protein synthesis of A549 cells after treatment with 25 VL/mL BMVs or 1 µM Antimycin A (AA) for 30 min (+ 30 min pre-treatment). Data were obtained from 5 biological replicates. **(D)** Overview of the effects of PA14 BMVs on the host cell and their underlying signaling pathway. Vesicles specifically inhibit complex III of the ETC resulting in mitochondrial dysfunction which activates the AMPK signaling pathway leading to an AMPK-dependent inhibition of protein synthesis via the translation regulator eEF2. Mitochondrial dysfunction and AMPK activation both induce a metabolic reprogramming of the host cell. Created with BioRender.com. All bar plots in this Figure are depicted as mean (*±* SEM). All significance niveaus were determined by Student’s t-test (** = p < 0.01).

As mentioned, AMPK is known to sense mitochondrial dysfunction caused by ETC inhibition. Toyama et al. have already shown that Antimycin A, a commonly used inhibitor of complex III of the ETC, can rapidly activate AMPK, leading to mitochondrial fragmentation [44]. A similar connection between mitochondrial dysfunction and AMPK activation is observed with the common type 2 diabetes drug metformin. Metformin inhibits complex I of the ETC and activates AMPK signaling, resulting in decreased cell proliferation and protein synthesis [50].

Since BMVs and Antimycin A both target ETC complex III, we wanted to test whether this impairment results in an inhibition of host cell protein synthesis. To that end, we treated A549 cells with PA14 BMVs and Antimycin A and analyzed global protein synthesis. We observed a similar inhibitory effect on global protein synthesis for both treatments (Figure 5C).

Taken together, our results suggest that the recognition of PA14 BMVs is not a direct process, mediated by cellular receptors, but occurs through mitochondrial dysfunction due to the specific inhibition of ETC complex III, which then activates AMPK signaling (Figure 5D).

In this study, we investigated the effects of *P. aeruginosa* PA14 BMVs on cell metabolism and mitochondrial respiration in human lung cells. In summary, we observed cellular metabolic reprogramming following vesicle treatment, particularly affecting TCA cycle-associated metabolites. In agreement with recent investigations [11], we identified vesicle-driven mitochondrial dysfunction, but more specifically, inhibition of complex III of the mitochondrial ETC by PA14 BMVs. We further demonstrated the activation of the metabolic sensor AMPK, presumably as a consequence of impaired complex III function. In conclusion, we identified AMPK-dependent inhibition of global protein synthesis in the host cell (Figure 5D).

