## Supplementary information for "Bacterial membrane vesicles of *Pseudomonas Aeruginosa* activate AMPK signaling through inhibition of mitochondrial complex III"

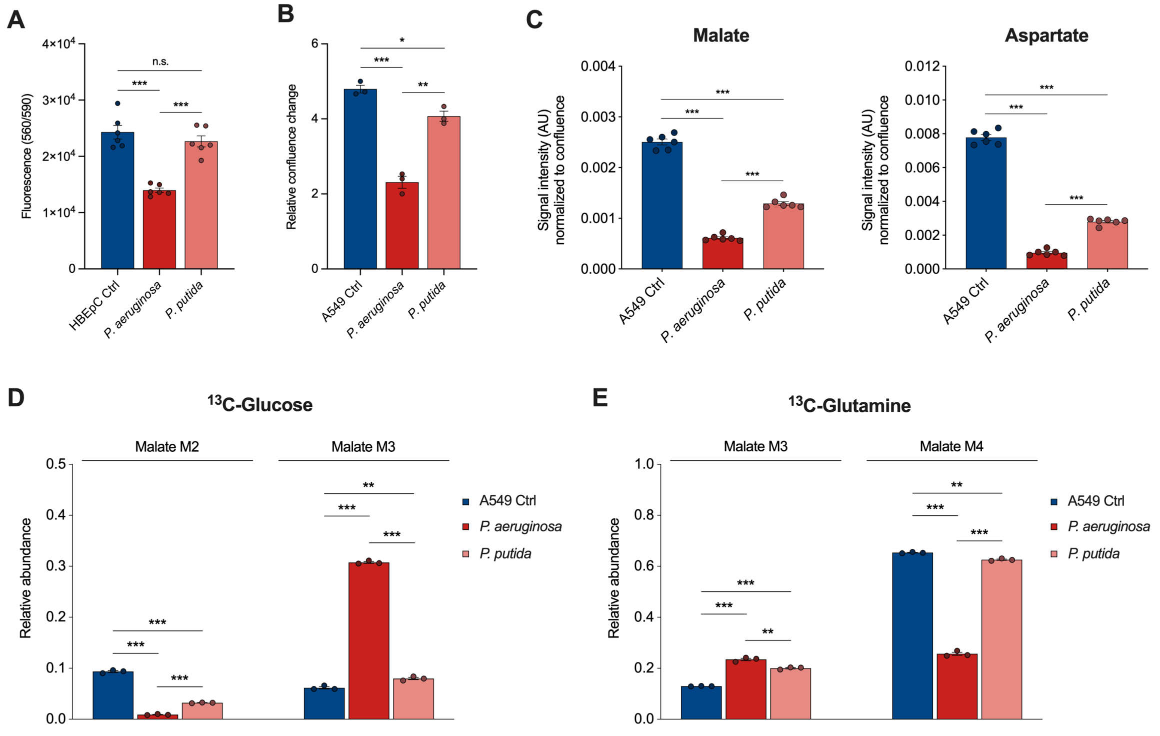


**Figure S1: Reduced effects of BMVs isolated from the non-pathogenic strain *P. putida* KT2400 compared to *P. aeruginosa* PA14 BMVs.** **(A)** Cell viability of HBEpC cells after treatment with 25 VL/mL BMVs for 24 h. Data were obtained from 6 biological replicates. **(B)** Confluence of A549 cells after treatment with 25 VL/mL BMVs for 72 h. Data were obtained from 3 biological replicates. **(C)** Signal intensities (AU) of malate and aspartate in A549 cells after vesicle treatment (25 VL/mL for 24 h). Data were obtained from 6 biological replicates and normalized to cell confluence. **(D)** Malate MIDs after [U-$^{13}C_6$]-glucose labeling of BMV-treated A549 cells (25 VL/mL for 24 h). Data were obtained from 3 biological replicates. **(E)** Malate MIDs after [U-$^{13}C_5$]-glutamine labeling of BMV-treated A549 cells (25 VL/mL for 24 h). Data were obtained from 3 biological replicates. All bar plots in this Figure are depicted as mean $\pm$ SEM. Significance niveaus were determined by Student's t-test (n.s. = not significant, * = p < 0.05, ** = p < 0.01, *** = p < 0.001).

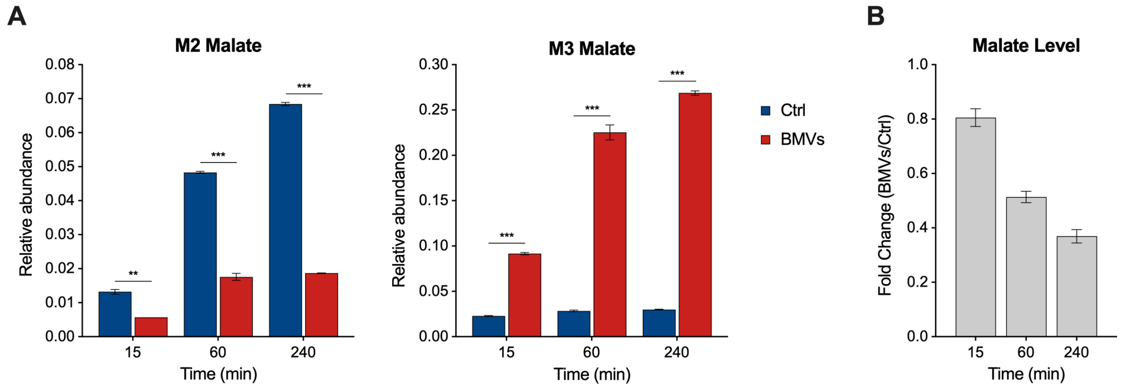


**Figure S2: Metabolic changes of A549 cells treated with PA14 BMVs after different timepoints. (A)** Malate MIDs of [U-$^{13}C_6$]-glucose labeled A549 cells after treatment with 25 VL/mL BMVs at three different timepoints (15, 60 and 240 min). Data were obtained from 2 or 3 biological replicates. **(B)** Relative signal intensities (fold change) of malate in BMV-treated A549 cells (25 VL/mL for 24 h) and cells without treatment (Ctrl). Data were obtained from 2 or 3 biological replicates. All bar plots in this Figure are depicted as mean $\pm$ SEM. Significance niveaus were determined by Student's t-test (** = p < 0.01, *** = p < 0.001).

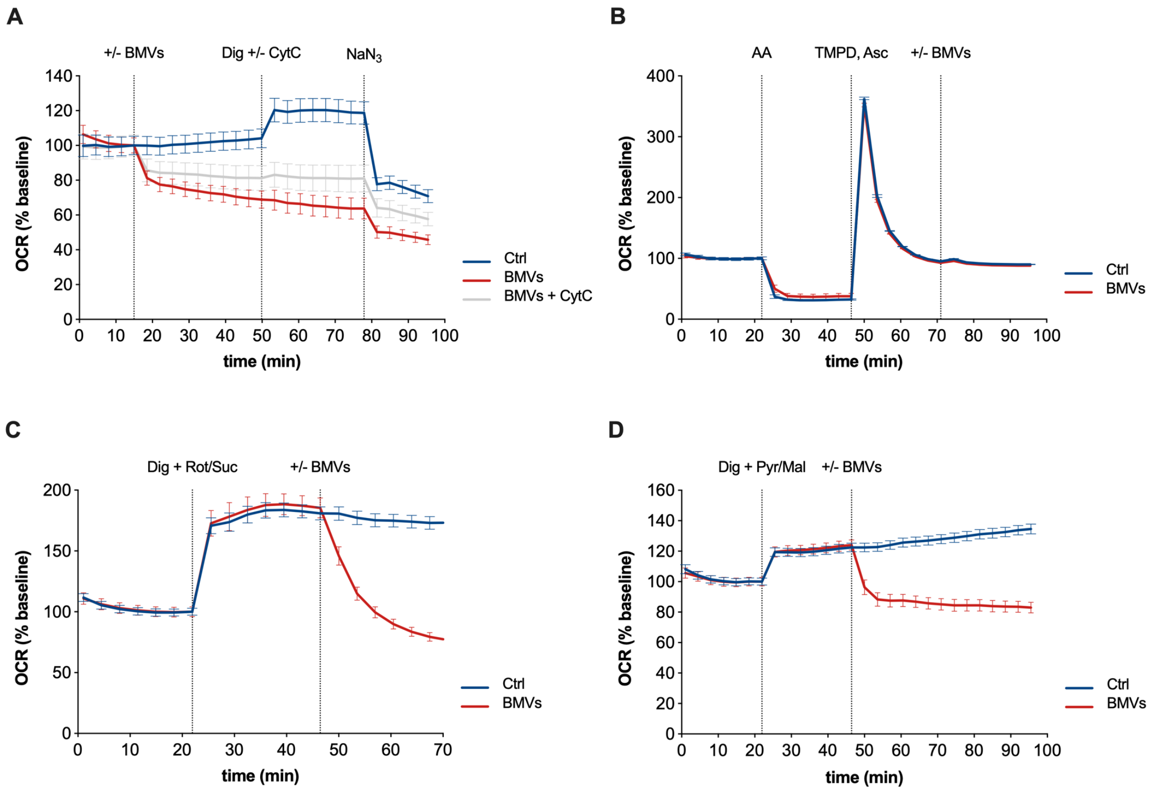


**Figure S3: Respirometry tests for the analysis of PA14 BMV-induced effects on the ETC of A549 cells.** **(A)** Respirometry test for Cytochrome *c* (CytC) activity of A549 cells treated with 25 VL/mL BMVs. Cytochrome *c* delivery (100 µM, blue and gray line) to mitochondria was accomplished by addition of 25 µg/mL digitonin (Dig), together with 5 mM succinate, 1 mM ADP, 5 mM EDTA and 3.6 mM phosphate ($K_2HPO_4$). For complex IV inhibition, 5 mM sodium azide ($NaN_3$) was added. Data were obtained from 20 biological replicates. **(B)** Respirometry test for complex IV activity of A549 cells treated with 25 VL/mL BMVs. Isolation of complex IV was performed by the addition of 0.5 µM Antimycin A (AA), 0.8 mM N,N,N',N'-tetramethyl-p-phenylenediamine (TMPD) and 16 mM ascorbate (Asc). Data were obtained from 15 biological replicates. **(C)** Respirometry test for complex II activity of A549 cells treated with 25 VL/mL BMVs. Isolation of complex II was performed by the addition of 25 µg/mL digitonin (Dig), together with the substrates rotenone (0.5 µM) and succinate (5 mM), 0.3 mM ADP, 5 mM EDTA and 3.6 mM phosphate ($K_2HPO_4$). Data were obtained from 15 biological replicates. **(D)** Respirometry test for complex I activity of A549 cells treated with 25 VL/mL BMVs. Isolation of complex I was performed by the addition of 25 µg/mL digitonin (Dig), together with the substrates pyruvate (5 mM) and malate (5 mM), 0.3 mM ADP, 5 mM EDTA and 3.6 mM phosphate ($K_2HPO_4$). Data were obtained from 15 biological replicates. OCR values were normalized to the baseline and depicted as mean $\pm$ SEM. Substances were added at the depicted time points.

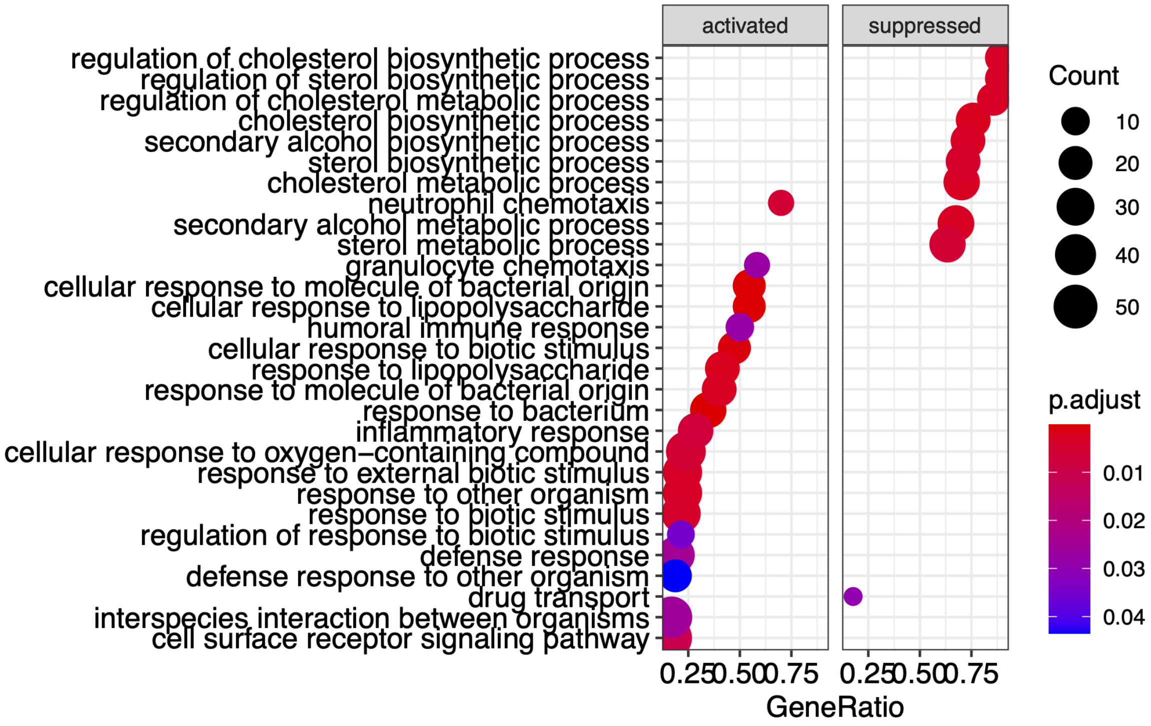


**Figure S4: Significantly enriched (activated and suppressed) GO-BP pathways in A549 cells following PA14 BMV treatment.** The vertical axis lists the names of GO-BP terms, while the length of the horizontal bars represents the gene ratio. The area of each circle indicates the gene counts, and the color intensity reflects the adjusted P-value. GO-BP: Gene Ontology biological process.

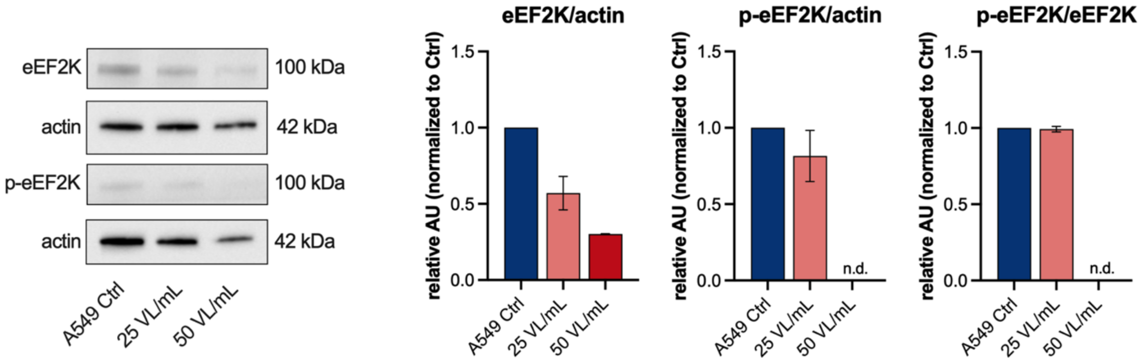


**Figure S5: Western blot analysis of the mTOR-specific phosphorylation site of eEF2K in A549 cells after PA14 BMV treatment.** WB analysis of p-eEF2K (Ser366) in A549 cells after vesicle treatment (25 VL/mL and 50 VL/mL) for 24 h. Data were obtained from 2 independent experiments. All bar plots in this Figure are depicted as mean (± SEM). n.d. = not detectable.
